# Purkinje cell collaterals preferentially target a subtype of molecular layer interneuron

**DOI:** 10.1101/2025.07.17.665322

**Authors:** EP Lackey, A Norton, L Moreira, C Gaynor, WA Lee, WG Regehr

## Abstract

In addition to providing outputs from the cerebellar cortex, Purkinje cell (PC) axon collaterals target other PCs, molecular layer interneurons (MLIs), and Purkinje layer interneurons (PLIs). It was recently shown that MLIs consist of two subtypes, but the properties of PC synapses onto these subtypes was not known and it was assumed that all PC collateral to MLI synapses would provide positive feedback to PCs. Clarifying the PC connectivity onto MLI subtypes is vital to understating the influence of feedback from PC collaterals because MLI1s primarily inhibit PCs whereas MLI2s mainly inhibit MLI1s and disinhibit PCs. Here we use a combination of serial EM and optogenetic studies to characterize PC synapses onto MLI subtypes in mice. EM reconstructions show that PCs make 53% of their synapses onto other PCs, 32% onto PLIs, 6% onto MLI1s and 7% onto MLI2s. Since there are far more MLI1s than MLI2s, each MLI2 is expected to receive many more synapses than each MLI1. In slice experiments, optogenetic activation of PCs evokes inhibitory currents in most MLI2s, but primarily disinhibits MLI1s. We also find that candelabrum cells, a type of PLI, form many more synapses onto MLI1s than MLI2s. It is therefore expected that both PC-MLI2-MLI1-PC and PC-PLI-MLI1-PC pathways allow increased PC firing to disinhibit MLI1s, which are known to reduce dendritic PC calcium signals and suppress plasticity at granule cell to PC synapses. These pathways provide negative feedback that act in concert with PC-PC synapses to counter elevations in PC firing.

**Significance Statement:** Purkinje cells (PCs) influence processing by inhibiting neurons in the cerebellar cortex, including other PCs, molecular layer interneurons (MLIs) and Purkinje layer interneurons (PLIs). The influence of PC-MLI synapses is not known because there are recently identified MLI subtypes with opposing effects: MLI1s inhibit PCs whereas MLI2s inhibit MLI1s and disinhibit PCs. We used serial EM and optogenetic studies to characterize PC synapses onto MLI subtypes and found that PCs preferentially inhibit MLI2s and disinhibit MLI1s. We also found that candelabrum cells (a type of PLI) preferentially inhibit MLI1s. These findings establish that PC-PC synapses, the PC-MLI2-MLI1-PC pathway and the PC-candelabrum cell-MLI1-PC pathway act together to allow alterations in PC firing to provide negative feedback to other PCs.

## Introduction

Purkinje cells (PCs) process tens of thousands of granule cell inputs and in turn provide outputs from the cerebellar cortex to the deep cerebellar nuclei, the vestibular nuclei and the parabrachial nucleus (Voogd and Glickstein 1998, Barmack 2003, Hashimoto et al. 2018, Chen et al. 2023, Chen et al. 2025). PC axons also have collaterals and synapse onto most types of neurons in the cerebellar cortex (Larramendi and Lemkey-Johnston 1970, Bishop 1988, O’Donoghue and Bishop 1990, Bernard et al. 1993, Bishop et al. 1993, Orduz and Llano 2007, Witter et al. 2016, Halverson et al. 2022). PCs inhibit other PCs, thereby allowing increases in PC firing to suppress the firing of other PCs (Orduz and Llano 2007, Witter et al. 2016). PC collaterals make many synapses near the PC layer and strongly inhibit the three types of Purkinje layer interneurons (PLIs): Lugaro cells, globular cells and candelabrum cells (Hirono et al. 2012, Witter et al. 2016, Miyazaki et al. 2021, Osorno et al. 2022). PC collaterals also extend into the molecular layer and inhibit approximately 25% of molecular layer interneurons (MLIs) (Witter et al. 2016). MLIs are potentially important targets because they are prevalent, with approximately 11 times as many MLIs as PCs (Korbo et al. 1993, Zhang et al. 2023) and 18 times as many MLIs as PLIs (Kozareva et al. 2021, Osorno et al. 2022). MLIs control PC firing, regulate calcium signaling in PC dendrites, and gate plasticity at granule cell to PC synapses (Dizon and Khodakhah 2011, Gaffield et al. 2018, Rowan et al. 2018, Kim and Augustine 2021). PCs even inhibit the input layer of some regions of the cerebellar cortex by directly synapsing onto granule cells and unipolar brush cells (UBCs) (Guo et al. 2016, Guo et al. 2021). In some cases, PCs exhibit target specificity, as for UBCs where they preferentially target ON UBCs (Guo et al. 2016, Guo et al. 2021). Conversely, PCs largely avoid some types of neurons, as is the case for Golgi cells that do not appear to be inhibited by collaterals (Witter et al. 2016).

Although much is known about PC collateral feedback, and it had long been thought that PCs inhibit MLIs (O’Donoghue et al. 1989, Bishop 1993, Bishop et al. 1993, Witter et al. 2016, Halverson et al. 2022), the recent discovery of two types of MLIs (Kozareva et al. 2021, Lackey et al. 2024) raises fundamental questions regarding the overall influence of PC collaterals. These two types of MLIs, MLI1 and MLI2, have very different effects on the cerebellar circuit. MLI1s are electrically coupled, they fire synchronously with each other, primarily inhibit PCs, suppress calcium signals in PC dendrites, and suppress plasticity at granule cell to PC synapses (Kozareva et al. 2021, Lackey et al. 2024). MLI2s are not electrically coupled, they primarily inhibit MLI1s and disinhibit PCs, and they are expected to promote plasticity at granule cell to PC synapses (Kozareva et al. 2021, Lackey et al. 2024). Previous studies of PC to MLI feedback were performed prior to the realization that there are two types of MLIs, and it was therefore assumed that PC inhibition of MLIs will in turn disinhibit PCs (Witter et al. 2016, Halverson et al. 2022). This would only be the case if PCs primarily inhibit MLI1s, whereas PC-MLI2 synapses are expected to suppress MLI1 firing and provide positive feedback by disinhibiting PCs. It is not known if PCs preferentially target one subtype of MLI. Similarly, the effects of PC-PLI feedback is unclear because it is not known if PLIs primarily inhibit MLI1s or MLI2s.

Here we assess the targets of PC collaterals using EM reconstructions, and recordings of optogenetically-evoked PC IPSCs onto identified MLI subtypes. EM reconstructions show that PCs make putative synapses onto PCs, PLIs and both MLI subtypes. There are slightly more putative synapses onto MLI2s than MLI1s, but it is expected that PCs more strongly inhibit each MLI2 because there are many more MLI1s than MLI2s (Kozareva et al. 2021, Lackey et al. 2024). Optogenetic activation of PCs evoked IPSCs in two thirds of MLI2s, but primarily disinhibited MLI1s. This disinhibition likely arises from a combination of PCs suppressing MLI2 and PLI firing that both in turn inhibit MLI1s. These findings suggest that PC collateral activation of MLI2s, PLIs and other PCs all provide negative feedback to PCs.

## Materials and Methods

### Slice experiments

Animal procedures were performed in accordance with the NIH and Animal Care and Use Committee (IACUC) guidelines and protocols approved by the Harvard Medical School Standing Committee on Animals. Pcp2-Cre Jdhu (B6.Cg-Tg(Pcp2-Cre)3555Jdhu/J) and ChR2-EYFP (B6;129S-Gt(ROSA)26Sor^tm32(CAG-COP4*H134R/EYFP)Hze^) mice were obtained from Jackson Laboratories. The Pcp2-Cre Jdhu line is selective for PCs (Witter et al. 2016). Animals of either sex were randomly selected for experiments. Animals were housed on a normal light–dark cycle with an ambient temperature of 18–23 °C with 40–60% humidity.

Acute parasagittal slices (220-μm thick) were prepared from P28-45 Pcp2-Cre Jdhu x ChR2-EYFP mice. Mice were anaesthetized with an intraperitoneal injection of ketamine (10 mg kg^−1^) and perfused transcardially with an ice-cold solution containing (in mM): 110 choline chloride, 7 MgCl_2_, 2.5 KCl, 1.25 NaH_2_PO_4_, 0.5 CaCl_2_, 25 glucose, 11.6 sodium ascorbate, 3.1 sodium pyruvate, 25 NaHCO_3_, equilibrated with 95% O_2_ and 5% CO_2_. Slices were cut in the same solution, and then transferred to artificial cerebrospinal fluid (ACSF) containing (in mM) 125 NaCl, 26 NaHCO_3_, 1.25 NaH_2_PO_4_, 2.5 KCl, 1 MgCl_2_, 1.5 CaCl_2_ and 25 glucose equilibrated with 95% O_2_ and 5% CO_2_. Following incubation at 34 °C for 30 min, the slices were kept up to 6 h at room temperature until recording.

Recordings were performed at 32 °C with an internal solution containing (in mM): 150 K-gluconate, 3 KCl, 10 HEPES, 3 MgATP, 0.5 GTP, 0.5 EGTA, 5 phosphocreatine-tris_2_ and 5 phosphocreatine-Na_2_ (pH adjusted to 7.2 with KOH, osmolarity adjusted to 310 mOsm kg^−1^). Visually guided whole-cell recordings were obtained with patch pipettes of ∼3-6-MΩ resistance pulled from borosilicate capillary glass (BF150-86-10, Sutter Instrument). Electrophysiology data were acquired using a Multiclamp 700A or 700B amplifier (Axon Instruments), digitized at 20 kHz and filtered at 4 kHz. Acquisition and analysis of slice electrophysiological data were performed using custom routines written in Igor Pro (Wavemetrics) and Matlab. (R)-CPP (2.5 µM) and NBQX (5 µM) were included in the bath solution to block glutamatergic receptors. All drugs were purchased from Abcam and Tocris.

Slices were kept in the dark before recording. Recordings were made from lobules IV-V of the vermis. We recorded MLIs in voltage clamp. We recorded from MLIs in the inner two-thirds of the molecular layer and determined the identity of MLI1s and MLI2s by characterizing their characteristic electrical properties as previously described (Kozareva et al. 2021, Lackey et al. 2024). We used spikelets to identify MLI1s, and a lack of spikelets combined with a high input resistance to identify MLI2s.

IPSCs were recorded at a holding potential of -45 mV. To activate PC boutons, we delivered brief pulses (1 ms) of blue light (∼60 mW/mm^2^) to a small spot (diameter ∼100 µm) at 1 Hz for 100 trials. The light stimulus was delivered using a laser (473 nm, Optoengine) or an LED (470 nm, Thorlabs), coupled through the excitation pathway of a BX50WI upright microscope (Olympus), and focused onto slices through a 40x or 60x water-immersion objective (Olympus).

Postsynaptic currents were low-pass filtered at 1 kHz and time-locked to the onset of stimulation. All synaptic currents are averages of 100 trials. The amplitudes of outward and inward currents were measured as the average of 3-5 ms (t_1_) and 15-20 ms (t_2_) following stimulation, respectively. Synaptic currents and z-score amplitudes were measured relative to baseline averaged 100 ms prior to the stimulation, and amplitudes at baseline were measured over t_1_ and t_2_ from 50 ms prior to stimulation. Based on the baseline measurements, responses to stimulation were determined to be significant if they had a z-score>2 for t_1_, and z-score<-2 for t_2_.

### Serial EM reconstructions

We previously imaged and aligned a 770 μm X 750 μm X 53 μm volume of lobule V of the mouse cerebellum for EM reconstructions comprised of 1176 45-nm thick parasagittal sections (Osorno et al. 2022, Nguyen et al. 2023, Lackey et al. 2024). We used automated image segmentation to generate neuron boundaries and automated synapse prediction to infer synaptic connectivity as described previously (Nguyen et al. 2023). We used the neuron segmentation to reconstruct PC axon collaterals and analyze their synaptic outputs. PCs are readily identified based on their distinctive large cell bodies and their extensive spiny dendrites. To rapidly reconstruct and correct errors from automated segmentation, we used a proofreading workflow called Dahlia, where annotators checked all neurons for split or merge errors. Synapses made by collaterals were identified using automated synapse detection as previously described (Nguyen et al. 2023). Synapses were proofread manually to validate each postsynaptic target using MD-Seg. Synapses were identified by characteristic ultrastructural features of GABAergic synapses, including a synaptic cleft with a flattening of opposed pre- and post-synaptic membranes and clustering of synaptic vesicles near the presynaptic specialization (Peters et al. 1991). Any automated detected synapses that did not fit these criteria were excluded from further analysis. By restricting our analysis six PC collaterals located near the center of the volume it was possible to identify almost all of their targets.

We identified MLIs targeted by PC collaterals and CCs as MLI1s or MLI2s based on their connectivity and morphology as previously described (Lackey et al. 2024). It was difficult to categorize many of the PLI targets into PLI subtypes based on morphological reconstructions from a 50 μm thick volume, because PLIs are less restricted to the parasagittal plane than MLIs. We therefore did not subdivide PLI targets into subtypes in our EM analysis with the exception of four candelabrum cells that we identified. Candelabrum cells were characterized previously based on the location of their cell body within the Purkinje cell layer, their pear shaped soma and beaded axons.

## Results

### EM reconstructions of PC collaterals and their targets

We initially characterize the connectivity of PC collaterals using a large-scale electron microscopy (EM) dataset and serial reconstructions from lobule V of a mouse cerebellum (Osorno et al. 2022, Nguyen et al. 2023, Lackey et al. 2024). Different types of neurons were identified based on their morphologies. The EM volume we study is approximately 50 μm thick and is well suited to capturing the dendritic and axonal arbors of neurons such as MLIs whose dendrites are parasagittal to the surface of the slice. PCs are readily identified based on their distinctive large cell bodies and their extensive spiny dendrites. In many cases PC axons reach the surface of the volume and are cut off, thereby preventing reconstructions of their collaterals and their targets. In lobule V, PC collaterals make most of their synapses near the PC layer and sometimes extend into the lower part of the molecular layer (Witter et al. 2016). Virtually all neurons in the molecular layer are MLIs. MLI1 and MLI2 subtypes were identified on the basis of their distinctive connectivity patterns, with MLI1s primarily contacting PCs and MLI2s mainly contacting MLI1s (Lackey et al. 2024). In addition, MLI1s in the lower two thirds of the molecular layer all contribute to a specialized structure known as a pinceaux near the initial segment of PC axons, but MLI2s do not contribute to pinceaux (Lackey et al. 2024). Within 40 μm of the bottom of the PC layer almost all MLIs are MLI1s (Lackey et al. 2024, Wang, 2022 #1341), whereas MLI1s and MLI2s are interspersed in the rest of the molecular layer. There are 3 times as many MLI1s as MLI2s in this data set (Lackey et al. 2024), which is consistent with the MLI1:MLI2 ratio in _sn_RNA_seq_ studies (Kozareva et al. 2021). However, for MLIs in the lower 60% of the molecular layer, whose dendrites are more likely to overlap with PC collaterals, the MLI1:MLI2 ratio is approximately 5:1. Golgi cells, whose cell bodies are located in the granular layer, have extensive dendrites that are often cut off. The cell bodies of PLIs are often located in the PC layer, but can be located in the lower molecular layer or the upper granular layer (Hirono et al. 2017, Miyazaki et al. 2021, Osorno et al. 2022). Their dendrites and axons are also often cut off. In some cases, PC collaterals contact unidentified pieces of dendrites that could correspond to MLI1s, MLI2s, PLIs or GCs, but by restricting our analysis to collaterals that are near the center of the volume it is possible to identify almost all of their targets.

As expected (Witter et al. 2016), reconstructed collaterals made extensive contacts near the PC layer, but also extended into the molecular layer (**Fig. 1a**). This collateral contacts multiple cell types including PCs (*grey*), MLI1s (*purple*) and MLI2s (*green*). An expanded view of the 3D reconstruction shows the PC synaptic contact onto an MLI2 **(Fig. 1b**, *top*) and the synapse is shown in a single section **(Fig. 1b**, *bottom*). A PC-MLI1 synapse (**Fig. 1c**) and a PC synapse onto another PC (**Fig. 1d**) are also shown. We previously used EM reconstructions to show that PCs collaterals also make putative synapses onto PLIs, specifically candelabrum cells (Osorno et al. 2022).

**Fig. 1.**
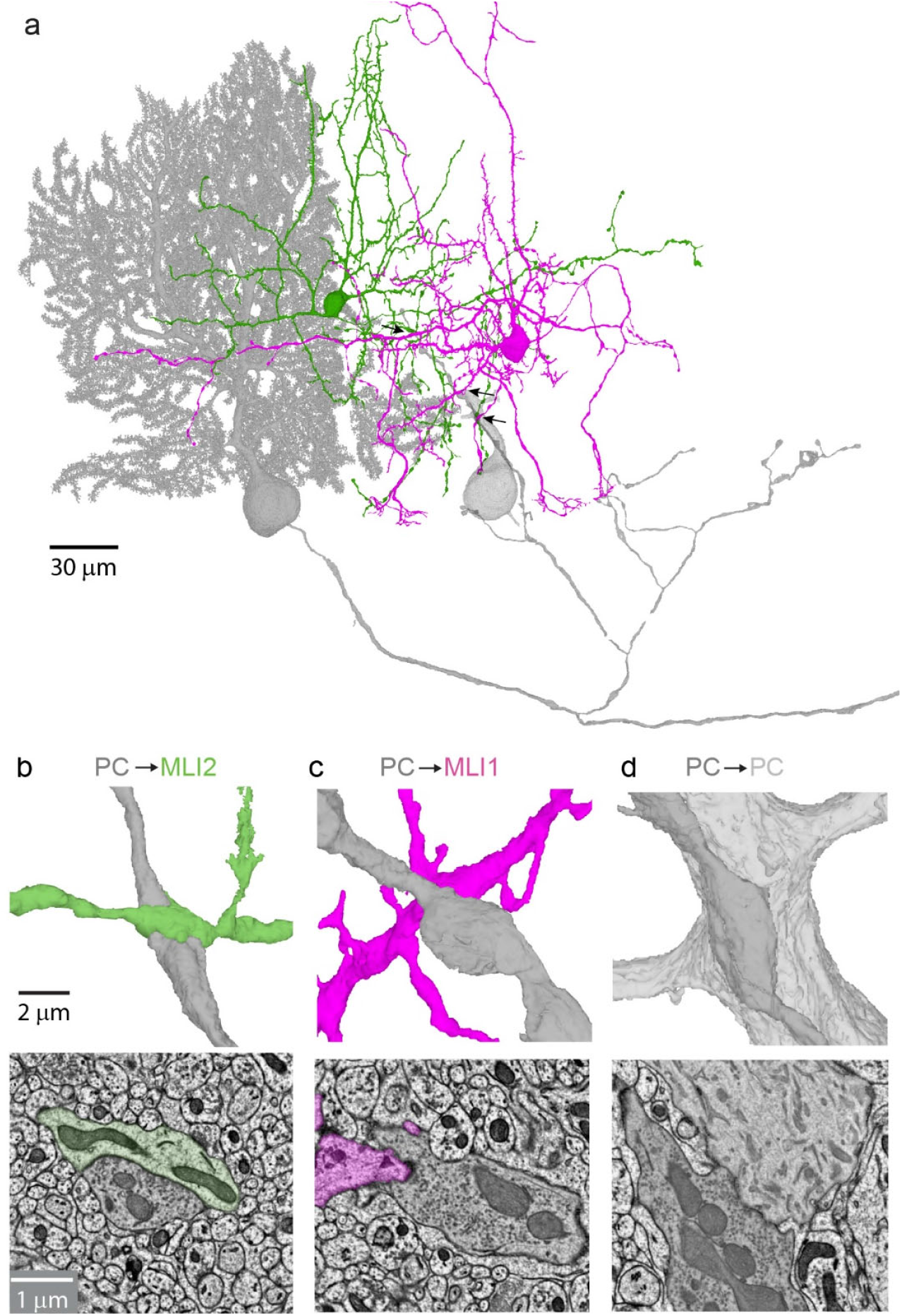
EM reconstructions reveals that PC collaterals target multiple cell types. **a**. Reconstructed PC and its collateral (*grey*), are shown along with 3 of its targets: an MLI2 (*green*), an MLI1 (*purple*), and another PC (*light grey*). Arrows indicate the locations of insets in **b**-**d**. **b**. Example of a PC-MLI1 connection shown with an expanded view of the 3D reconstruction (*top*) and with a single section (*bottom*). **c**. As in **b** but for a PC-MLI2 connection. **d**. As in **b** but for a PC-PC connection.

We reconstructed 6 collaterals, identified 244 synaptic boutons, and determined the identities of all but one of the targets (**Fig. 2**). Synaptic contacts were made onto PCs, PLIs and MLIs. There were no synaptic contacts onto Golgi cells, which is consistent with previous optogenetic studies (Witter et al. 2016). It was possible to determine the subtype identity of the MLIs, but it was difficult to determine the specific PLI subtypes based on morphological reconstructions from a 50 μm thick volume. We therefore did not subdivide PLIs into subtypes (although see candelabrum cells in **Fig. 4**). On average, each collateral made 40.5 ± 5.6 putative synapses (range 28 - 61). There was considerable variability in the targets of different collaterals (**Fig. 2a**)..The percentage of synapses onto different targets was 53.1 ± 4.0 % (range 42 - 60 %) onto PCs, 35.8 ± .3 % (range 20-51 %) onto PLIs, 4.8± 1.5 % (range 0-7 %) onto MLI1s, and 5.7 ± 2.8 % (range 0 - 17 %) onto MLI2s (**Fig. 2b**).

**Fig. 2.**
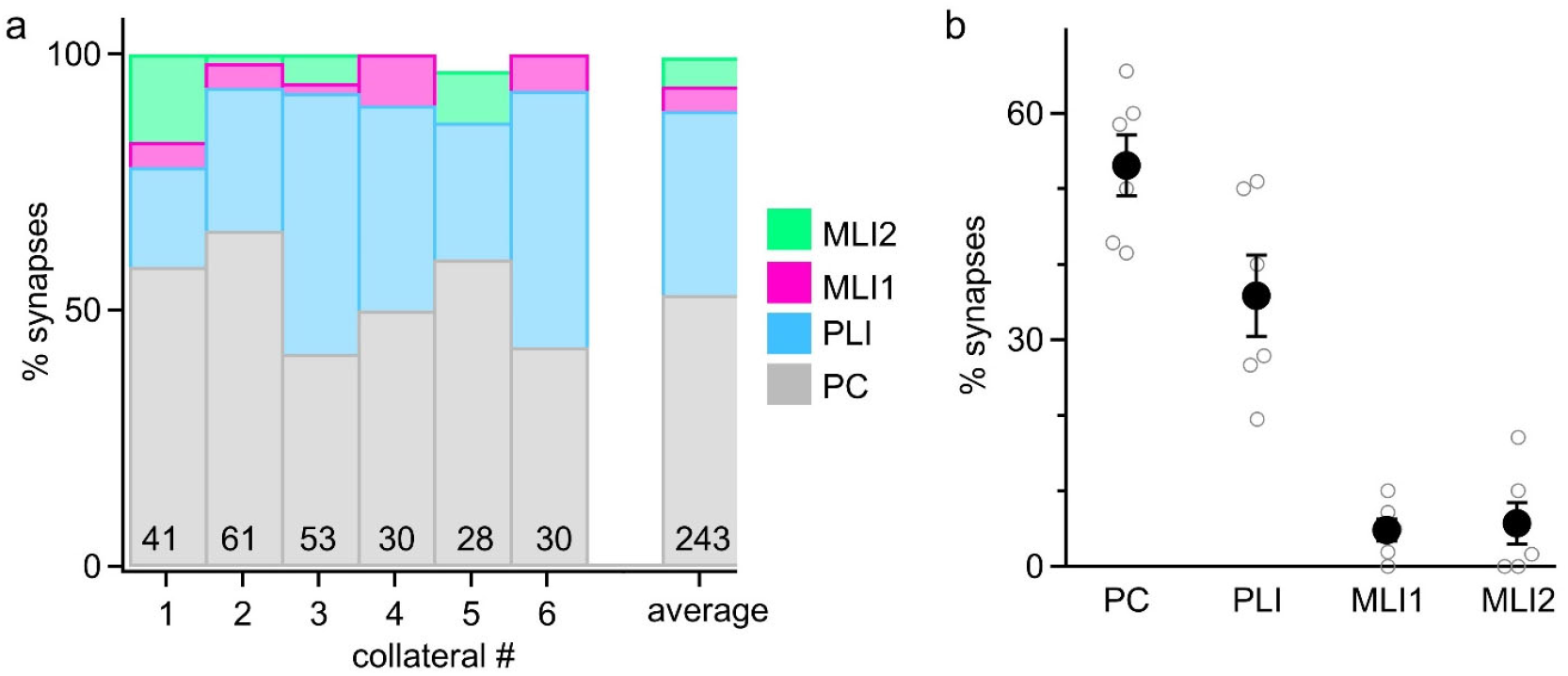
Summary of reconstructions of PC collaterals and their targets. **a**. Summary of the percentages of PC collateral synapses onto different targets. The total numbers of putative synapses for each collateral are indicated. The average for all collaterals is also shown. **b**. Percentages of synapses made onto different types of targets by different collaterals.

It is possible to use these data to estimate the relative number of putative synaptic contacts onto MLI1s, and MLI2s, which are present at a ratio of approximately 5:1 for cells in the inner 60% of the molecular layer. Our reconstructions indicate that there are on average approximately 5.9 (5*5.7/4.8) times as many putative PC contacts onto MLI2s as MLI1s and suggests that on average PC-MLI2 synapses are likely to be much stronger than PC-MLI1 synapses.

### Optogenetically-evoked PC-MLI Synapses

In addition to the number of putative synapses, which can be readily estimated from serial reconstructions, factors such as the probability of release, postsynaptic receptors and quantal size can influence synaptic strength. We therefore used an optogenetic approach to study PC-MLI synapses in brain slices (**Fig. 3a**). Experiments were performed using Pcp2-Cre Jdhu mice to selectively target PCs (Zhang et al. 2004) that were crossed with a line to conditionally express ChR2-EYFP in PCs (Witter et al. 2016). Recordings were made from MLI subtypes that were distinguished on the basis of their electrical properties, including their resistance and the presence of spikelets that are a hallmark of electrically-coupled cells (Alcami and Marty 2013). One of the most striking differences between MLI subtypes is that MLI1s are electrically coupled to each other, whereas MLI2s are not (Kozareva et al. 2021, Lackey et al. 2024). The spontaneous firing of a neighboring electrically coupled MLI1 gives rise to spikelets in nearby MLI1s that are a distinguishing feature of MLI1s (Kozareva et al. 2021, Lackey et al. 2024). Spikelets are not observed in MLI2s. Recordings were restricted to MLIs in the low two thirds of the molecular layer where electrical coupling between MLI1s is stronger and differences between MLI1 and MLI2 input resistances are larger (Kozareva et al. 2021, Lackey et al. 2024). In addition, collaterals are more prominent in the lower part of the molecular layer (Witter et al. 2016). The ratio of the number of MLI1s to the number of MLI2s in our recordings was 4.9:1 (44 MLI1s and 9 MLI2s), which is consistent with the known distribution of MLI subtypes within 120 μm of the bottom of the PC layer.

**Fig. 3.**
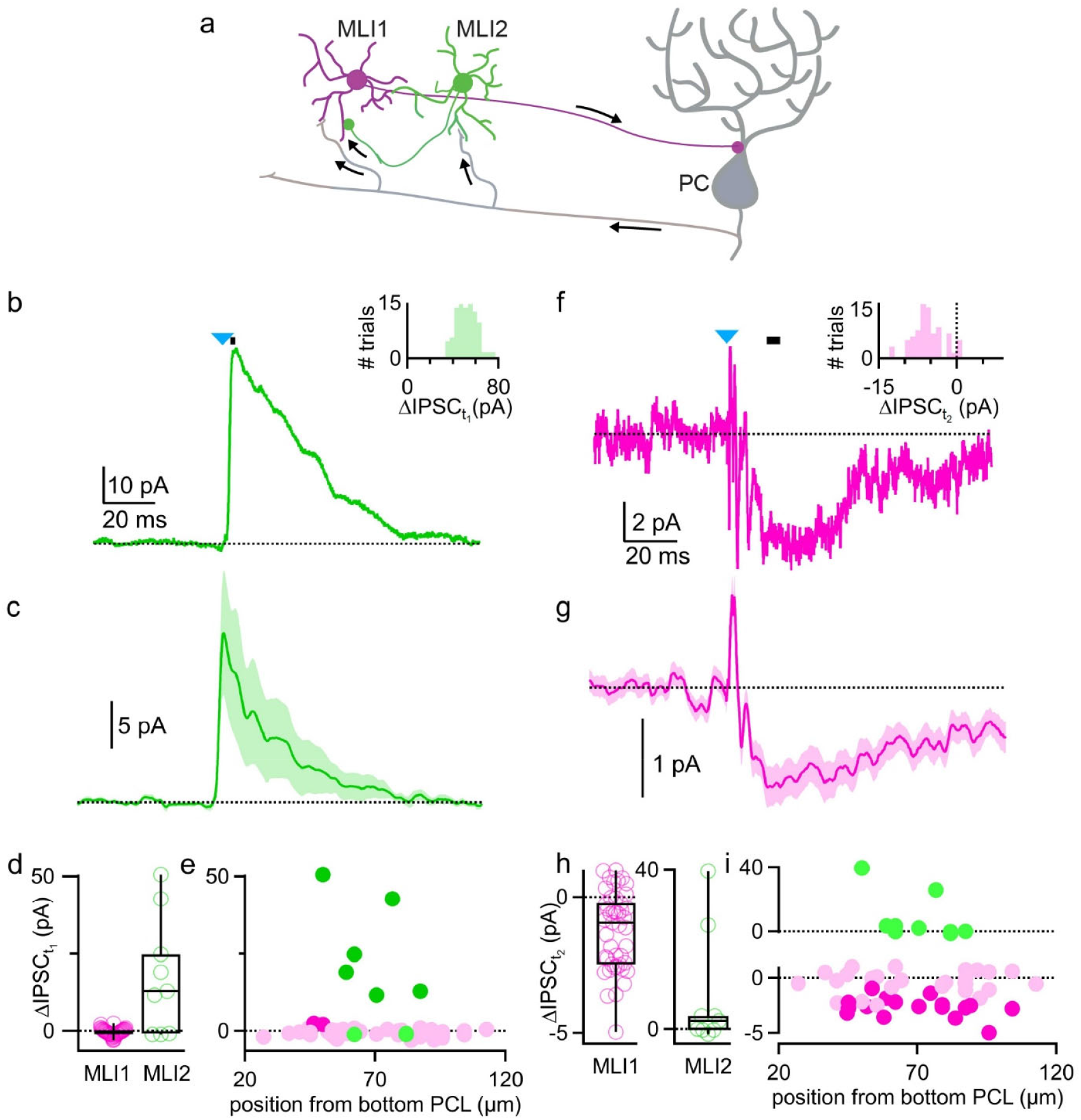
Optogenetic activation of PCs preferentially inhibits MLI2s. **a**. Schematic of slice experiments in which PCs in PC-ChR2 mice were activated with blue light and whole-cell recordings were made from MLI1s and MLI2s. **b**. Light-activated synaptic response in an MLI2. Inset: The histogram of shows the amplitudes of 100 responses for this cell. Black line indicates 3-5 ms following light activation. The blue triangle indicates the timing of optogenetic activation. **c**. Average ± SEM of synaptic responses evoked in 9 MLI2s. **d**. Summary of the optical synaptic responses evoked in MLI1s and MLI2s (averages of 3-5 ms following light activation). **e**. Amplitudes of evoked responses 3-5 ms following optical stimulation for individual cells as a function of distance from the bottom of the PCL layer. Significant responses were evoked in six MLI2s (*bold green*) and responses in the remaining MLI2s (*faint green*) and most of the MLI1s (*faint purple*) were not significant. Small but significant inhibition was observed in 2 of 44 MLI1s. **f**. Light-activated synaptic response in an MLI1. Inset: The histogram of shows the amplitudes of 100 responses for this cell. Black line indicates 15-20 ms following light activation. **g**. Average ± SEM of synaptic responses evoked in 9 MLI2s. **h**. Summary of the optical synaptic responses evoked in MLI1s and MLI2s (averages of 15-20 ms following light activation). **i**. Amplitudes of evoked responses 15-20 ms following optical stimulation for individual cells as a function of distance from the bottom of the PCL layer. Significant responses were evoked in 17 MLI1s (dark purple) and responses in the remaining MLI1s (*faint purple*) and all of the MLI2s (*faint green*) were not significantly decreased.

Optogenetic activation of PCs evoked synaptic responses in most MLI2s (**Fig. 3b-d**). As shown for an example MLI2, these responses could be large and reliable (**Fig. 3b**). In this case, optogenetic activation always evoked a response and there were never any failures, as shown by a histogram of the amplitudes of responses evoked in 400 trials (**Fig. 3b**, *inset*). For the cells with PC-MLI2 connections, failures were observed in 22 ± 9% (n=6) of the trials. The example optical PC-MLI2 IPSC was quite slow, and had a half decay time of 29 ms (**Fig. 3c**). The prolonged time course of this response likely reflects the ability of a single brief (0.5 ms) optical stimulus to evoked multiple spikes in PCs. Optically-evoked responses were detected in 6 of 9 MLI2s located between 50 μm and 87 μm from the bottom of the PC layer (**Fig. 3d**).

Light-evoked responses were very different in neurons identified on the basis of spikelets and low input resistances. An extremely large IPSC (61 pA) was evoked in 1 of 45 neurons that satisfied this criteria (**Extended data Fig. 1**), with the rest of the neurons having average responses near 0 (-3.0 to 2.6, n=44) and weak but significant PC-MLI1 inhibition observed in 2 of 44 cells. We therefore did not categorize this neuron as an MLI1 because it was an outlier with properties more consistent with a displaced PLI, which are not strictly located within the PC layer (Osorno et al. 2022). For the remaining 44 neurons that we categorized as MLI1s, rather than evoking an inhibitory IPSC, light activation evoked an inward current in many of these neurons, as in **Fig. 3e**. These responses were small (**Fig. 3f**) and highly variable, as shown by a histogram of the magnitude of suppression for 100 trials (**Fig. 3f**, inset). The variability is not surprising considering that the responses are tiny, and there are many spontaneous IPSCs that introduce noise. The net inward current is consistent with PC inhibition of MLI2s leading to disinhibition of MLI1s. The average evoked responses from ML1s were very small and quite slow (**Fig. 3g**). Disinhibition was seen in 38% (17 of 45 with a Z-score of < -2) of MLI1s that were located between 45 μm and 104 μm from the bottom of the PC layer (**Fig. 3hi**).

### Identifying MLI subtypes targeted by candelabrum cells

To understand how the PC to PLI synapses influence processing in the molecular layer, it is necessary to determine which MLI subtypes are targeted by PLIs. Unfortunately, the axons of most PLIs were not confined to the volume and it was not possible to reconstruct them sufficiently to determine their synaptic targets. However, we previously reconstructed PC synapses onto candelabrum cells and found that candelabrum cells made extensive synapses onto MLIs, but the identity of MLI targets was not known (Osorno et al. 2022). We therefore used serial EM to identity the MLI subtypes targeted by candelabrum cells (**Fig. 4**). In **Fig. 4a**, the cell body and the axon of the same PC as in **Fig. 1** is shown (grey) along with a candelabrum cell that it targets (*blue*). An expanded view shows a PC synapse onto this candelabrum cell (**Fig. 4b**). Although the dendrite of this candelabrum cell is incomplete, its distinctive beaded axon extends from the PC layer to the top of the molecular layer where it forms synapses onto 22 MLI1s (*purple*) and 6 MLI2s (*green*). Candelabrum to MLI2 (**Fig. 4c**) and candelabrum to MLI1 (**Fig. 4d**) synapses are shown in expanded views. We found that four candelabrum cells (whose axons were reconstructed with different degrees of completeness) preferentially targeted MLI1s: candelabrum cells made 249 synapses onto MLI1s and 39 synapses onto MLI2s, so that for PC-MLI synapses, 86.5% are onto MLI1s and 13.5% are onto MLI2s (**Fig. 4e**).

**Fig. 4.**
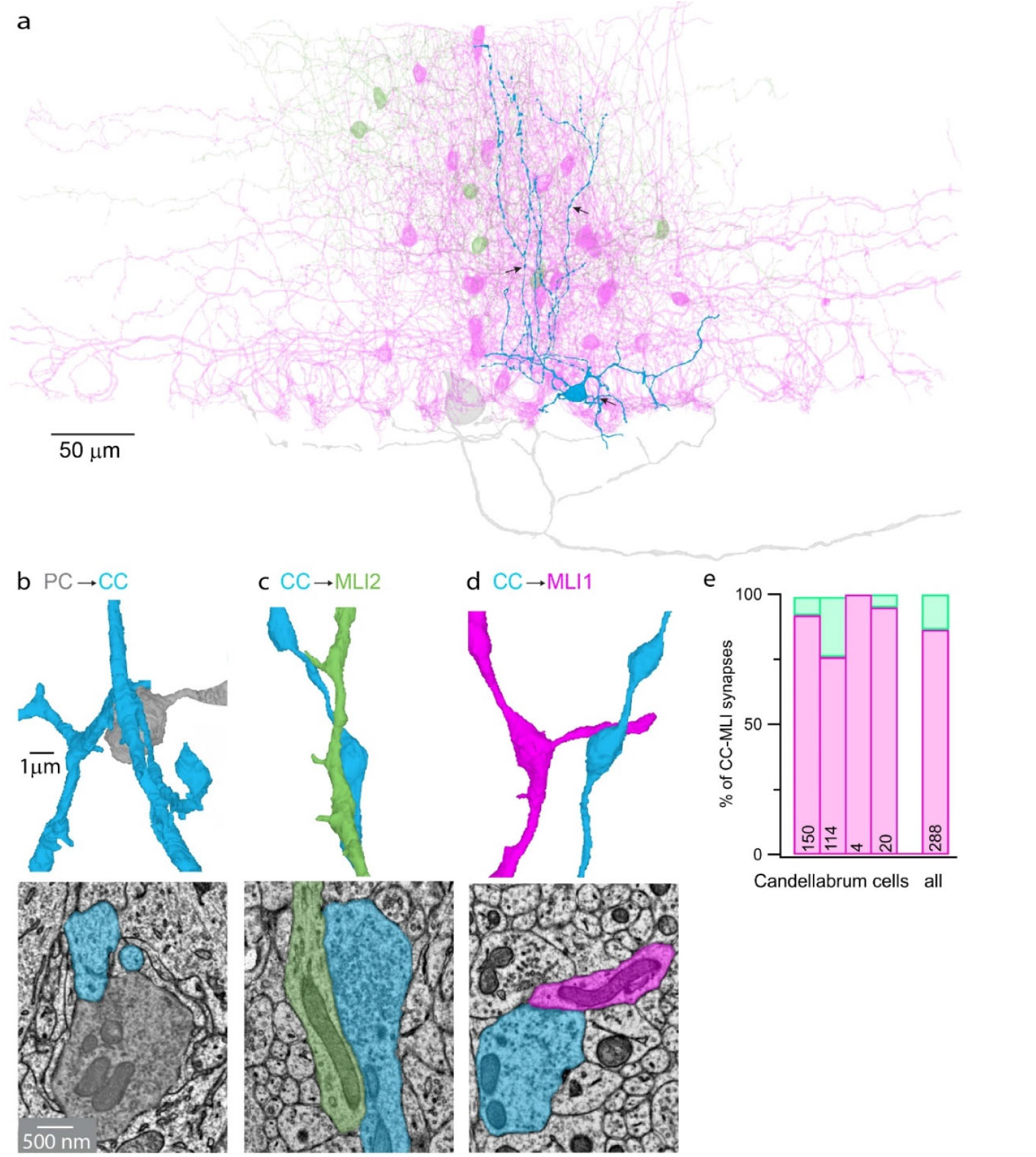
Candelabrum cells preferentially target MLI1s. **a**. 3D EM reconstruction of MLI1s (*purple*) and MLI2s (*green*) targeted by a candelabrum cell (*blue)* that receives inhibitory input from a PC collateral (*grey*). Inset shows the synapses onto MLI1s and MLI2s. The dendrites of the candelabrum cell are incomplete because they are cut off by the surface of the slice. The PC dendrite is not shown for clarity. Arrows indicate the locations of regions shown in **b**-**d**. **b**. Example of a PC-candelabrum cell connection shown with an expanded view of the 3D reconstruction (*top*) and with a single section (*bottom*). **c**. As in **b** but for a candelabrum cell to MLI2 synapse. **d**. As in **b** but for a candelabrum cell to MLI1 synapse. **e**. Summary of candelabrum cell synapses onto MLI1s and MLI2s for 4 collaterals, with the number of reconstructed synapses indicted. The total number of synapses onto different MLI subtypes is also shown to the right. This is an extension of previous reconstructions with the exception that the identities of the MLI subtypes had not been determined (Osorno et al. 2022).

## Discussion

Our main finding is that PCs selectively activate MLI2s and primarily disinhibit MLI1s. This allows PC feedback to MLI2s to ultimately suppress PC firing. This is a major revision of the view of PC to MLI feedback, which was thought to lead to disinhibition of PCs prior to the recent identification of MLI subtypes. Moreover, our findings establish that there are multiple pathways that work in concert to provide negative feedback to PCs and suppress PC firing: the PC-MLI2-MLI1-PC, PC-candelabrum cell-MLI1-PC, and PC-PC pathways (**Fig. 5**).

**Fig. 5.**
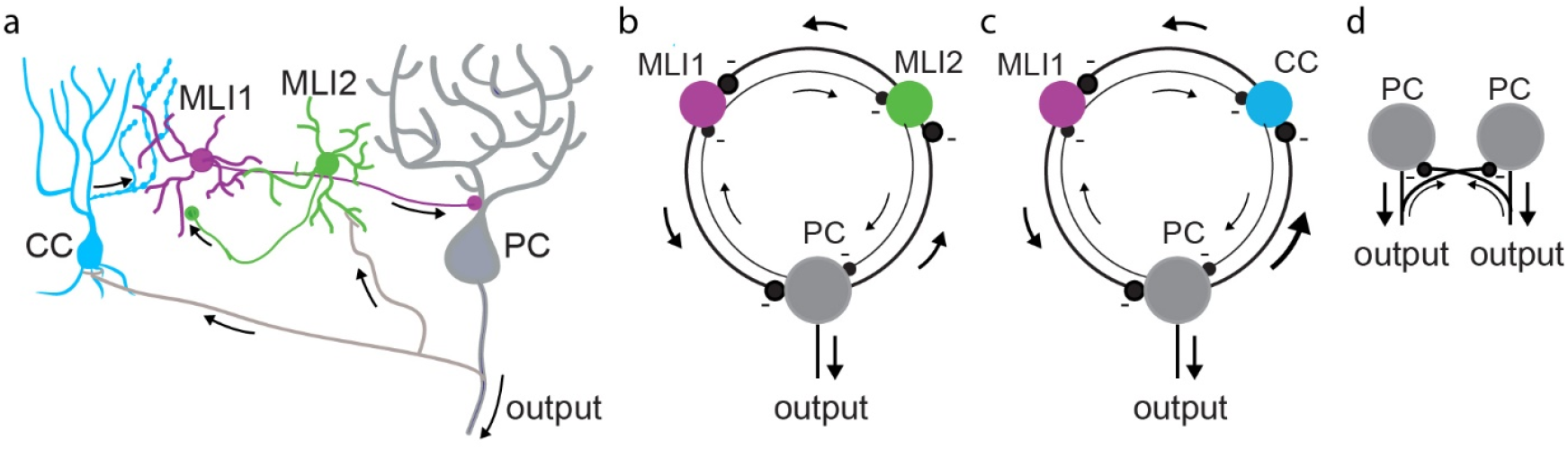
Circuit diagrams of PC collaterals and their interneuron targets in the cerebellar cortex. a. Schematics of PC collaterals and their targets, MLI1s, MLI2s and candelabrum cells (CCs). b. Three-ring inhibitory loop involving PCs, MLI2s, and MLI1s. c. Three-ring inhibitory loop involving PCs, CCs, and MLI1s. d. Two-ring inhibitory loop involving PCs.

### PC synapses preferentially target MLI2s

EM reconstructions and slice electrophysiology both indicate that PCs preferentially inhibit MLI2s. Reconstructions indicate that although PC collaterals make a comparable number of putative synaptic contacts onto MLI1s and MLI2s, the approximately 5-fold difference in the number of MLI1s within 120 μm of the PC layer suggests that each MLI2 will receive PC inputs that are approximately 6 (5*5.7/4.8) times larger. Optogenetic studies were also consistent with a much stronger PC inhibition of MLI2s than MLI1s: two thirds of MLI2s were directly inhibited by PCs and inhibitory PC-MLI1 connections were rarely observed. Based on EM reconstructions, we were surprised that only very weak PC-MLI1 inhibition was observed in 2 of 44 cells. It is possible that PC-MLI2-MLI1 and PC-PLI-MLI1 disinhibition obscures small direct PC-MLI1 inhibition, or that each PC-MLI1 synapse is weaker than each PC-MLI2 because their probability of release is lower, or they are associated with a lower number of GABA_A_ receptors. Our results indicate that PCs do not provide prominent inhibition to MLI1s and but lead to MLI1 disinhibition and are part of a PC-MLI2-MLI1-PC loop (**Fig. 5b**)

### The PC-PLI-MLI pathway

Our reconstructions also provide insight into the functional role of PC to PLI feedback (**Fig. 5c**). Without knowing subtype specificity of PLI-MLI synapses, it was not known if this pathway promotes plasticity and elevates PC firing or suppresses plasticity and reduces PC firing. We were able to address this issue for candelabrum cells, which are the most numerous subtype of PLI, and our reconstructions established that candelabrum cells make 5.9 times as many synapses onto MLI1s than onto MLI2s (**Fig. 4**). In contrast to PC collaterals that are restricted to the PC layer and the lower part of the molecular layer, candelabrum cell axons extend to the surface of the molecular layer and form synapses throughout this region (Laine and Axelrad 1994, Osorno et al. 2022). Therefore, candelabrum cell synapses are in the vicinity of 3 times as many MLI1s as MLI2s, and based solely on EM reconstructions we estimate that on average candelabrum cell synapses are approximately twice (5.9/3) as strong onto each MLI1s than each MLI2s. The PC-candelabrum cell-MLI circuit is therefore well suited to allowing elevated PC activity to disinhibit MLI1s. These findings are consistent with the observation that candelabrum cell activation suppresses spontaneous IPSCs in PCs (Osorno et al. 2022). Although it is well established that PCs also strongly inhibit Lugaro cells and globular cells, we were unable to reconstruct the axons of these cells and determine which subtype of MLI they preferentially target. Based on the molecular similarity of Lugaro cells, globular cells and candelabrum cells, and to a somewhat lesser extent MLI2s (Kozareva et al. 2021), it seems likely that Lugaro cells and globular cells also preferentially target MLI1s, but further experiments are needed to determine if this is the case.

### Overview of Targets of PC Collaterals

The characterization of all synapses made by PC collaterals provides an overview of the relative strengths of different pathways. PC collateral synapses are made onto 3 types of targets: 53% PC, 36% PLI, and 11% MLI. These findings are consistent with optogenetic studies in which light-evoked PC IPSCs were large and consistently evoked in PCs and Lugaro cells (a type of PLI), but they were only observed in 25% of the MLIs (Witter et al. 2016). Consistent with our finding that PCs selectively activate MLI2s, this percentage of MLIs is comparable to the percentage of MLI2s in our snRNAseq study (24.5%, 10,608 MLI2s / 43,324 total MLIs) (Kozareva et al. 2021), and the percentage of MLI2s in our EM dataset (24%, 26 MLI2s / 108 total MLIs) (Lackey et al. 2024). Although the PC-PC, PC-MLI2-MLI1-PC and PC-candelabrum cell-MLI1 pathways all allow elevated PC firing to provide negative feedback and suppress PC firing (**Fig. 5**), it is likely that there are important differences in their contributions to cerebellar processing.

### PC Collaterals and Plasticity

It is likely that PC collateral inhibition of other PCs and collateral pathways that lead to disinhibition of MLIs differentially regulates long-term depression (LTD) of granule cell to PC synapses. When PC firing rates are elevated, the PC-MLI2-MLI1-PC and PC-candelabrum cell-MLI1-PC pathways will increase MLI1 firing frequency, reduce climbing fiber induced calcium signals in PC dendrites and suppress LTD (Callaway et al. 1995, Rowan et al. 2018). In contrast the PC-PC pathway only leads to activation of inhibitory synapses onto the proximal PC dendrite and the soma and is not expected to exert a strong influence on dendritic calcium signals and LTD. Thus, although all three pathways are expected to provide negative feedback to PCs and suppress firing, activation of MLI2s and candelabrum cells are expected to be much more effective at regulating dendritic calcium and plasticity.

### PC Collaterals and Oscillations

Different PC collateral pathways likely contribute to oscillatory PC firing in different ways. PCs can generate 150-200 Hz oscillations in mice that can be time locked to movements (Adrian 1935, Dow 1938, Isope et al. 2002, de Solages et al. 2008, Groth and Sahin 2015, Cheron et al. 2016). It is thought that PC to PC inhibition (**Fig. 5d**) underlies these oscillations. Mutual inhibition of PCs is well suited to generate oscillatory firing at a frequency determined by the synaptic delay and the time constant of the PC (Orduz and Llano 2007, de Solages et al. 2008). The PC-MLI2-MLI1-PC and PC-candelabrum cell-MLI1-PC loops are also suited to generating oscillations. These circuits have the architecture of ring oscillators, which consist of an odd number of inverting stages connected in a ring (Still and Le Masson 1999, Horikawa 2011). This circuit generates oscillations with a frequency (*f*) given by *f*=(1/2*tn*), where *n* is the number of stages and *t* is the delay at each stage (Razavi 2017). Digital electronics exploits this property and generates oscillations using ring oscillators (Razavi 2017). For loops of inhibitory neurons in the cerebellum, delays are the time taken to suppress firing in target cells, which is determined by the synaptic delay and time constants of the different cells. Loops involving MLIs are unlikely to contribute to the 150 to 200 Hz oscillations because: (1) we estimate that the delays are more consistent with generating slower oscillations in the 30 to 100 Hz range, (2) 150 to 200 Hz oscillations are amplified by activation of cannabinoid CB1Rs that strongly suppresses MLI to PC synapses, but leaves PC-PC synapses intact (de Solages et al. 2008), and (3) the power spectra of the MLI spike-triggered average does not have the 200 Hz peak that is prominent in the PC spike triggered average (Blot et al. 2016). It is possible that loops involving MLIs do not produce ongoing oscillatory activity, but that they provide transient or context-dependent negative feedback. Further experimental work will be needed to clarify the conditions under which these loops influence PC spiking.

## Acknowledgments

This work was supported by the NIH (R35NS097284 to W.G.R., F32NS133036 to E.P.L., R21NS085320, and RF1MH128949, and RF1MH114047 to W.-C.A.L.), and the Edward R. and Anne G. Lefler Center. Portions of this research were conducted on the O2 High Performance Compute Cluster at Harvard Medical School.

## References

Adrian, E. (1935). “Discharge frequencies in the cerebral and cerebellar cortex. .” Proc. Phys. Soc. 83: 32–33.

Alcami, P. and A. Marty (2013). “Estimating functional connectivity in an electrically coupled interneuron network.” Proc Natl Acad Sci U S A 110(49): E4798–4807.

Barmack, N. H. (2003). “Central vestibular system: vestibular nuclei and posterior cerebellum.” Brain Res Bull 60(5-6): 511–541.

Bernard, C., H. Axelrad and B. Giraud (1993). “Effects of collateral inhibition in a model of the immature rat cerebellar cortex: multineuron correlations.” Cognitive brain research 1(2): 100–122.

Bishop, G. A. (1988). “Quantitative analysis of the recurrent collaterals derived from Purkinje cells in zone x of the cat’s vermis.” J Comp Neurol 274(1): 17–31.

Bishop, G. A. (1993). “An analysis of HRP-filled basket cell axons in the cat’s cerebellum. I. Morphometry and configuration.” Anat Embryol (Berl) 188(3): 287–297.

Bishop, G. A., Y. F. Chen, R. W. Burry and J. S. King (1993). “An analysis of GABAergic afferents to basket cell bodies in the cat’s cerebellum.” Brain Res 623(2): 293–298.

Blot, A., C. de Solages, S. Ostojic, G. Szapiro, V. Hakim and C. Lena (2016). “Time-invariant feed-forward inhibition of Purkinje cells in the cerebellar cortex in vivo.” J Physiol 594(10): 2729–2749.

Callaway, J. C., N. Lasser-Ross and W. N. Ross (1995). “IPSPs strongly inhibit climbing fiber-activated [Ca2+]i increases in the dendrites of cerebellar Purkinje neurons.” J Neurosci 15(4): 2777–2787.

Chen, C. H., L. N. Newman, A. P. Stark, K. E. Bond, D. Zhang, S. Nardone, C. R. Vanderburg, N. M. Nadaf, Z. Yao, K. Mutume, I. Flaquer, B. B. Lowell, E. Z. Macosko and W. G. Regehr (2023). “A Purkinje cell to parabrachial nucleus pathway enables broad cerebellar influence over the forebrain.” Nat Neurosci 26(11): 1929–1941.

Chen, C. H., Z. Yao, S. Wu and W. G. Regehr (2025). “Characterization of direct Purkinje cell outputs to the brainstem.” Elife 13.

Cheron, G., J. Marquez-Ruiz and B. Dan (2016). “Oscillations, Timing, Plasticity, and Learning in the Cerebellum.” Cerebellum 15(2): 122–138.

de Solages, C., G. Szapiro, N. Brunel, V. Hakim, P. Isope, P. Buisseret, C. Rousseau, B. Barbour and C. Lena (2008). “High-frequency organization and synchrony of activity in the purkinje cell layer of the cerebellum.” Neuron 58(5): 775–788.

Dizon, M. J. and K. Khodakhah (2011). “The role of interneurons in shaping Purkinje cell responses in the cerebellar cortex.” J Neurosci 31(29): 10463–10473.

Dow, R. S. (1938). “The electrical activity of the cerebellum and its functional significance.” J Physiol 94(1): 67–86.

Gaffield, M. A., M. J. M. Rowan, S. B. Amat, H. Hirai and J. M. Christie (2018). “Inhibition gates supralinear Ca(2+) signaling in Purkinje cell dendrites during practiced movements.” Elife 7.

Groth, J. D. and M. Sahin (2015). “High frequency synchrony in the cerebellar cortex during goal directed movements.” Front Syst Neurosci 9: 98.

Guo, C., S. Rudolph, M. E. Neuwirth and W. G. Regehr (2021). “Purkinje cell outputs selectively inhibit a subset of unipolar brush cells in the input layer of the cerebellar cortex.” Elife 10.

Guo, C., L. Witter, S. Rudolph, H. L. Elliott, K. A. Ennis and W. G. Regehr (2016). “Purkinje Cells Directly Inhibit Granule Cells in Specialized Regions of the Cerebellar Cortex.” Neuron 91(6): 1330–1341.

Halverson, H. E., J. Kim, A. Khilkevich, M. D. Mauk and G. J. Augustine (2022). “Feedback inhibition underlies new computational functions of cerebellar interneurons.” Elife 11.

Hashimoto, M., A. Yamanaka, S. Kato, M. Tanifuji, K. Kobayashi and H. Yaginuma (2018). “Anatomical Evidence for a Direct Projection from Purkinje Cells in the Mouse Cerebellar Vermis to Medial Parabrachial Nucleus.” Front Neural Circuits 12: 6.

Hirono, M., S. Nagao, Y. Yanagawa and S. Konishi (2017). “Monoaminergic modulation of GABAergic transmission onto cerebellar globular cells.” Neuropharmacology 118: 79–89.

Hirono, M., F. Saitow, M. Kudo, H. Suzuki, Y. Yanagawa, M. Yamada, S. Nagao, S. Konishi and K. Obata (2012). “Cerebellar globular cells receive monoaminergic excitation and monosynaptic inhibition from Purkinje cells.” PLoS One 7(1): e29663.

Horikawa, Y. (2011). “Exponential transient propagating oscillations in a ring of spiking neurons with unidirectional slow inhibitory synaptic coupling.” J Theor Biol 289: 151–159.

Isope, P., S. Dieudonne and B. Barbour (2002). “Temporal organization of activity in the cerebellar cortex: a manifesto for synchrony.” Ann N Y Acad Sci 978: 164–174.

Kim, J. and G. J. Augustine (2021). “Molecular Layer Interneurons: Key Elements of Cerebellar Network Computation and Behavior.” Neuroscience 462: 22–35.

Korbo, L., B. B. Andersen, O. Ladefoged and A. Moller (1993). “Total numbers of various cell types in rat cerebellar cortex estimated using an unbiased stereological method.” Brain Res 609(1-2): 262–268.

Kozareva, V., C. Martin, T. Osorno, S. Rudolph, C. Guo, C. Vanderburg, N. Nadaf, A. Regev, W. G. Regehr and E. Macosko (2021). “A transcriptomic atlas of mouse cerebellar cortex comprehensively defines cell types.” Nature 598(7879): 214–219.

Lackey, E. P., L. Moreira, A. Norton, M. E. Hemelt, T. Osorno, T. M. Nguyen, E. Z. Macosko, W. A. Lee, C. A. Hull and W. G. Regehr (2024). “Specialized connectivity of molecular layer interneuron subtypes leads to disinhibition and synchronous inhibition of cerebellar Purkinje cells.” Neuron.

Laine, J. and H. Axelrad (1994). “The candelabrum cell: a new interneuron in the cerebellar cortex.” J Comp Neurol 339(2): 159–173.

Larramendi, L. M. and N. Lemkey-Johnston (1970). “The distribution of recurrent Purkinje collateral synapses in the mouse cerebellar cortex: an electron microscopic study.” J Comp Neurol 138(4): 451–459.

Miyazaki, T., M. Yamasaki, K. F. Tanaka and M. Watanabe (2021). “Compartmentalized Input-Output Organization of Lugaro Cells in the Cerebellar Cortex.” Neuroscience 462: 89–105.

Nguyen, T. M., L. A. Thomas, J. L. Rhoades, I. Ricchi, X. C. Yuan, A. Sheridan, D. G. C. Hildebrand, J. Funke, W. G. Regehr and W. A. Lee (2023). “Structured cerebellar connectivity supports resilient pattern separation.” Nature 613(7944): 543–549.

O’Donoghue, D. L. and G. A. Bishop (1990). “A quantitative analysis of the distribution of Purkinje cell axonal collaterals in different zones of the cat’s cerebellum: an intracellular HRP study.” Exp Brain Res 80(1): 63–71.

O’Donoghue, D. L., J. S. King and G. A. Bishop (1989). “Physiological and anatomical studies of the interactions between Purkinje cells and basket cells in the cat’s cerebellar cortex: evidence for a unitary relationship.” J Neurosci 9(6): 2141–2150.

Orduz, D. and I. Llano (2007). “Recurrent axon collaterals underlie facilitating synapses between cerebellar Purkinje cells.” Proc Natl Acad Sci U S A 104(45): 17831–17836.

Osorno, T., S. Rudolph, T. Nguyen, V. Kozareva, N. M. Nadaf, A. Norton, E. Z. Macosko, W. A. Lee and W. G. Regehr (2022). “Candelabrum cells are ubiquitous cerebellar cortex interneurons with specialized circuit properties.” Nat Neurosci 25(6): 702–713.

Peters, A., S. L. Palay and H. d. Webster (1991). The fine structure of the nervous system : neurons and their supporting cells. New York, Oxford University Press.

Razavi, B. (2017). Design of Analog CMOS Integrated Circuits. New York, NY, McGraw-Hill.

Rowan, M. J. M., A. Bonnan, K. Zhang, S. B. Amat, C. Kikuchi, H. Taniguchi, G. J. Augustine and J. M. Christie (2018). “Graded Control of Climbing-Fiber-Mediated Plasticity and Learning by Inhibition in the Cerebellum.” Neuron 99(5): 999–1015 e1016.

Still, S. and G. Le Masson (1999). “Traveling waves in a ring of three inhibitory coupled model neurons.” Neurocomputing 26-27: 533–539.

Tukey, J. W. (1977). Exploratory Data Analysis, Addison-Wesley Publishing Company.

Voogd, J. and M. Glickstein (1998). “The anatomy of the cerebellum.” Trends Cogn Sci 2(9): 307–313.

Witter, L., S. Rudolph, R. T. Pressler, S. I. Lahlaf and W. G. Regehr (2016). “Purkinje Cell Collaterals Enable Output Signals from the Cerebellar Cortex to Feed Back to Purkinje Cells and Interneurons.” Neuron 91(2): 312–319.

Zhang, K., Z. Yang, M. A. Gaffield, G. G. Gross, D. B. Arnold and J. M. Christie (2023). “Molecular layer disinhibition unlocks climbing-fiber-instructed motor learning in the cerebellum.” bioRxiv: 2023.2008.2004.552059.

Zhang, X. M., A. H. Ng, J. A. Tanner, W. T. Wu, N. G. Copeland, N. A. Jenkins and J. D. Huang (2004). “Highly restricted expression of Cre recombinase in cerebellar Purkinje cells.” Genesis 40(1): 45–51.

