## Supplemental Figure 1 for "Purkinje cell collaterals preferentially target a subtype of molecular layer interneuron"

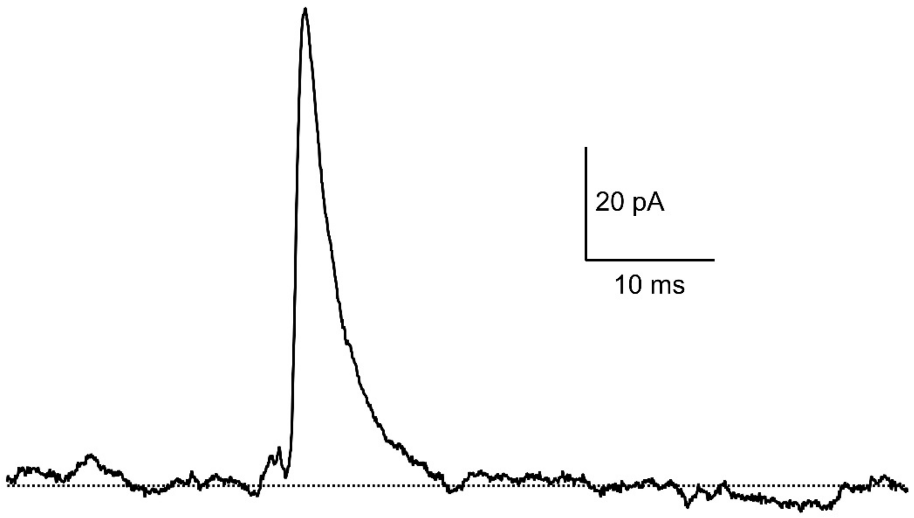


**Supplemental Fig. 1: Synaptic response from an outlier interneuron in the molecular layer.**

The synaptic response from a neuron located in the molecular layer 40 µm from the bottom of the PCL layer. The input resistance of the cell was 126 MΩ and spikelets were apparent. The amplitude of the optically-evoked IPSC was 61 pA, whereas all other responses in MLI1s were in the -3.0 to 2.6 pA range (n=44). Applying the interquartile range method (Tukey 1977) to the data set, all optically-evoked IPSCs are within the interquartile range of -3.4 to 3.0 pA, with the exception of this response which is well outside the interquartile range. The Grubbs test also indicates that it is an outlier (Grubbs statistic 6.53, t statistic 43, p value 3X $10^{-44}$). We therefore categorized this cell as an outlier and removed it from our data set. The large short-latency response suggests that this could be a displaced PLI.
